# PyBoost: A parallelized Python implementation of 2D boosting with hierarchies

**DOI:** 10.1101/170803

**Authors:** Peyton G. Greenside, Nadine Hussami, Jessica Chang, Anshul Kundaje

**Affiliations:** Biomedical Informatics, Stanford University, Stanford, 94305, USA; Electrical Engineering, Stanford University, Stanford, 94305, USA; Genetics, Stanford University, Stanford, 94305, USA; Computer Science, Stanford University, Stanford, 94305, USA

## Abstract

**Motivation:** Gene expression is controlled by networks of transcription factors that bind specific sequence motifs in regulatory DNA elements such as promoters and enhancers. GeneClass is a boosting-based algorithm that learns gene regulatory networks from complementary paired feature sets such as transcription factor expression levels and binding motifs across conditions. This algorithm can be used to predict functional genomics measures of cell state, such as gene expression and chromatin accessibility, in different cellular conditions. We present a parallelized, Python-based implementation of GeneClass, called PyBoost, along with a novel hierarchical implementation of the algorithm, called HiBoost. HiBoost allows regulatory logic to be constrained to a hierarchical group of conditions or cell types. The software can be used to dissect differentiation cascades, time courses or other perturbation data that naturally form a hierarchy or trajectory. We demonstrate the application of PyBoost and HiBoost to learn regulators of tadpole tail regeneration and hematopoeitic stem cell differentiation and validate learned regulators through an inducible CRISPR system.

**Availability:** The implementation is publicly available here: https://github.com/kundajelab/boosting2D/.

## 1 Introduction

Gene expression is carefully regulated by sets of transcription factor complexes binding specific DNA sequence motifs to translate the same genome into many different cell types from heart cells to brain cells. However, the details of each cell’s unique regulatory program remain poorly understood. To this end, many methods have been developed to decipher the regulatory mechanisms that control gene expression across different cell types and conditions [13] [2]. GeneClass and its sister algorithm MEDUSA, which incorporates motif discovery, are examples of such algorithms that have been successfully used to learn gene regulatory programs and predict gene expression dynamics across different conditions [6]. Recently, assays for chromatin accessibility such as DNase-seq and ATAC-seq have become popular for genome-wide characterization of cellular state. However, predicting chromatin accessibility genome-wide presents a substantially larger computational burden than predicting a fixed set of expressed genes. The original implementation of GeneClass, implemented in Matlab, is prohibitively slow for these higher dimensional applications. FastMEDUSA implements the MEDUSA algorithm in C++ to reduce the computational burden, but includes motif discovery, which may only apply to specific prediction problems, does not include postprocessing modules to interpret the models and may be harder to further develop for those not versed in C++ [1]. We introduce a parallelized, Python-based implementation of GeneClass, called PyBoost, to address this computational limitation and enable easy integration with postprocessing analysis. We provide accompanying analysis functionalities to determine key regulators in any set of conditions and for any set of regions or genes.

We also introduce a new implementation, called HiBoost, that enforces a hierarchy over conditions in the target matrix. Many experimental conditions have an inherent structure that can be defined hierarchically. For example, the differentiation of a stem cell into diverse cell types has a natural hierarchy starting at the stem cell root and cascading through intermediate states to the terminal cell types. Time courses also represent a natural order where each time step derives from a previous time step hierarchically. PyBoost learns weak classifiers that are maximally predictive over the entire data set. HiBoost allows each rule to be constrained to a specific part of the hierarchy placed upon the conditions to learn regulatory programs specific to that sub-tree of the hierarchy only. For example, in hematopoeisis certain regulatory rules may only apply to the myeloid lineage and not the lymphoid lineage and HiBoost enables this distinction. Our implementation allows improved interpretation of each regulatory rule in the context of the condition hierarchy and can substantially improve interpretability of the model overall by allowing context-specific rules.

## 2 Methods

### 2.1 Algorithm

#### 2.1.1 PyBoost

PyBoost, like GeneClass, is a boosting-based algorithm that builds Alternating Decisions Trees (ADTs) of weak classifiers. Weak classifiers or "rules" are composed of a paired motif and regulator, added on to an existing node in the ADT, that give a predicted classification. Training labels can be gene expression, chromatin accessibility or other phenotypes. The goal of pyBoost is to predict a binary output or target matrix *S* of dimension (element *e* x condition *c*) where an element can correspond to specific genes or transcripts, non-coding regions for accessibility, or other genomic measures. Conditions may be different cell types or experimental conditions. The paired feature sets required for this prediction, each corresponding to one dimension of the target matrix, are a binary matrix *M* of motif hits of dimension (motif *m* x element *e*) and a binary matrix *R* of regulator dynamics of dimension (condition *c* x regulator *r*). At each iteration, the motif and regulator with minimum loss are selected with a corresponding score α indicating the direction and strength of prediction. This rule can be added to any existing node in the ADT. After each iteration, the examples are re-weighted to prioritize those examples that have not been correctly predicted and the next rule is chosen based on this weighting of the training data. The result of this algorithm after many iterations is a gene regulatory network, in the form of an ADT, consisting of motif and regulator pairs with corresponding prediction scores and the sets of examples they apply to. Each of these rules contributes additively to the margin of prediction for each example, which can be analyzed after generating the ADT to understand how predictions were made. See [5] for further algorithm details.

#### 2.1.2 HiBoost

HiBoost extends the PyBoost algorithm to incorporate a hierarchy placed upon the conditions in the target matrix. Weak classifiers are extended from (motif *m*, regulator *r*, ADT node parent *p*, score *s*) to include the node in the condition hierarchy to which that specific rule applies, giving each rule the updated form (motif *m*, regulator *r*, ADT node parent *p*, score *s*, hierarchy node *h*). The rule is then applied to all conditions that fall within the subtree of the hierarchy rooted at the rule’s hierarchy node *h*. When adding new nodes to the ADT at each iteration, the new ADT node can be added at the current *h* of the node being added to or at any of the direct children of *h*, chosen by which of these nodes has the minimal loss at the given iteration. Thus each path down the ADT also represents a trajectory down the condition hierarchy [Figure 1].

**Fig. 1.**
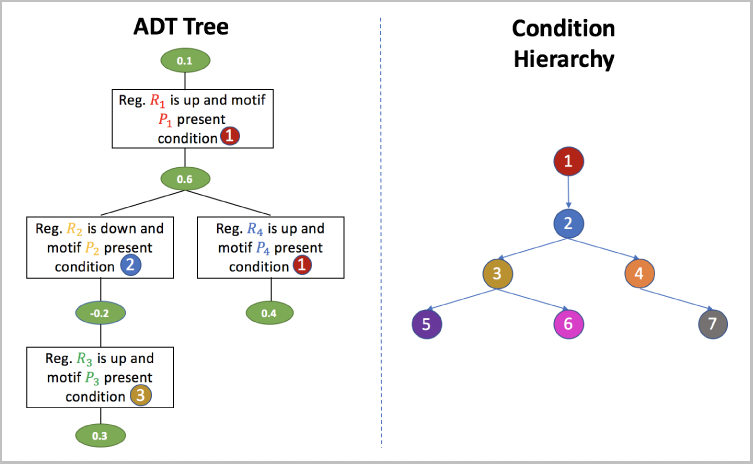
HiBoost extends the standard ADT from PyBoost by adding to each rule a node in the condition hierarchy such that the ADT node applies to the condition subtree rooted at that condition hierarchy node.

### 2.2 Implementation

PyBoost and HiBoost are implemented in Python 2.7 with all freely available libraries. The algorithm can be parallelized across a user-specified number of threads. The number of nodes to check for each new rule grows at every iteration and parallelization allows simultaneous evaluation of every current ADT node for the minimum loss. The algorithm can also run with sparse instantiations of feature and weight matrices to further reduce runtime. In addition to the prior, stabilization, and postprocessing modules discussed below, user inputs enable the generation of predictive stumps in place of an ADT structure, the ability to shuffle any of the three input matrices, which can be useful in identifying an empirical null model for a given ADT, the ability to supply a specified holdout matrix for train/validation splits, and an option to compress regulators that have the same pattern across conditions for improved interpretation.

#### 2.2.1 Prior Matrix

While the algorithm may pair any motif with any regulator for the best predictive power, many regulators are known to bind to particular types of motifs. To facilitate the association of regulators with biologically plausible motifs, we enable a prior matrix *P_mr_* over all motif-regulator pairs to prioritize meaningful pairings between motifs and regulators. In addition, we enable a prior matrix between regulators *P_rr_* to preferentially add rules on to ADT nodes where there is a known interaction between the current regulator and the regulator in the parent ADT node. The initial loss *L*_0_ for both priors is tuned by a prior constant *c*, indicating the strength of the prior, as well as a decay rate *d*, which can reduce the influence of the prior over many iterations, to generate the loss augmented by the prior: *L_P_* = *L*_0_(1+*Pcd^n^*). The decay rate can be used to prioritize establishing known regulatory programs in early nodes of the ADT before lessening the prior matrix to enable discovery of novel regulatory programs.

#### 2.2.2 Stabilization

Multiple motif-regulator pairs will often have highly similar or identical loss. As in Kundaje et al. [6], we enable stabilization of the algorithm by allowing multiple motif-regulator pairs that have loss within a small range of the absolute minimum loss to be included in a given rule. The corresponding rule bundle size can be tuned through user provided parameters. In HiBoost, all of the bundled rules are added at the same condition hierarchy node.

#### 2.2.3 Post-Processing

We provide margin-based post-processing tools to extract feature sets either motifs, regulators, ADT nodes or ADT paths - that regulate specific examples. For a user-provided set of conditions *c* and/or elements *e*, we rank feature sets by their contribution to the margin for that specified subset of the data. For making comparisons between features, margin scores for each feature are normalized by the number of examples they apply to. This tool can be used to track the relative importance of features across conditions or between regions. The example-by-feature matrix that outputs the normalized margin score for each example by each feature can be used in downstream analysis such as unsupervised clustering of feature sets or examples.

## 3 Results

### 3.1 PyBoost: Tadpole tail regeneration

We applied PyBoost to learn key regulators of tadpole tail regeneration. We profiled gene expression (RNA-Seq) and chromatin accessibility (ATACseq) at a series of time points during tail regeneration after the tail was injured. One of the top regulators discovered by PyBoost was SPIB. The influence of this regulator on tail regeneration was validated through an inducible CRISPR system [Figure 2].

**Fig. 2.**
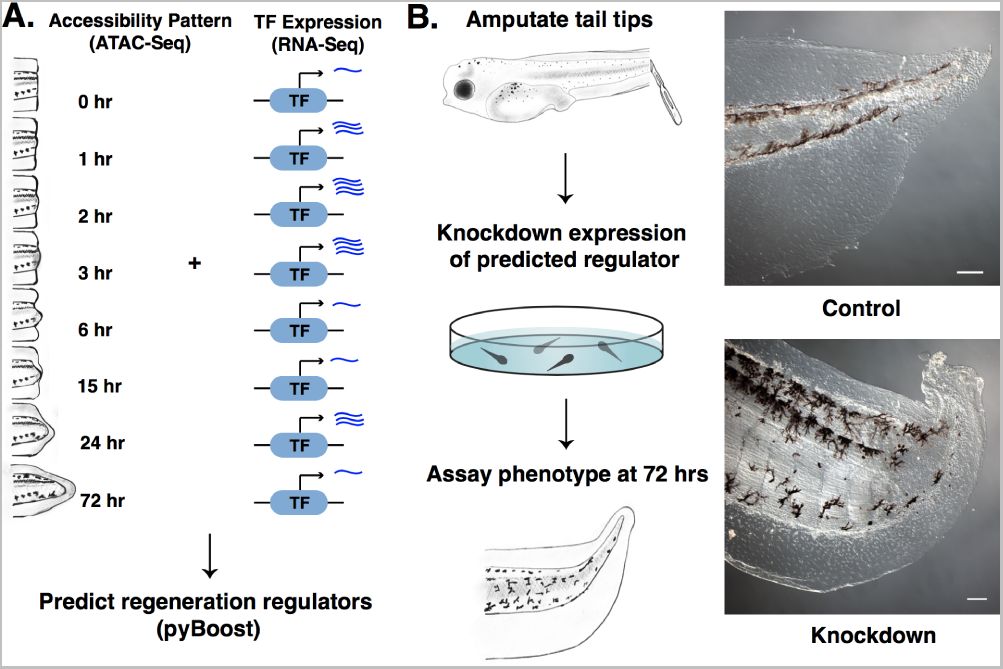
A. PyBoost was used to discover key regulators of tadpole tail regeneration from paired expression and accessibility data profiled in a time course of tail regeneration. B. SPIB was discovered as a top regulator and validated to strongly affect tail regeneration through an inducible CRISPR system.

### 3.2 HiBoost: Hematopoietic differentiation hierarchy

We applied HiBoost to learn regulatory programs governing chromatin accessibility dynamics during differentiation of hematopoieitic stems cells into 13 healthy terminal blood cells as well as 3 leukemic cell types [4]. HiBoost allows condition-specific rules at any point in the differentiation hierarchy. Out of 500 iterations, 278 were placed on the root hierarchical node while the remaining 222 were placed on non-root hierarchy nodes that would not have been available without the hierarchy implementation. The training and testing errors were similar between hierarchical (0.228 train error, 0.231 test error) and non-hierarchical methods (0.228 train error, 0.230 test error). Multiple rules were placed on the leukemic tree allowing context-specific interpretation of these regulators. The 7 unique proteins identified through these rules all have known leukemic associations including BACH1 [9], PBX1 [10], NFATC3 [8], ILKZF1 [11], MSX2 [14], HES1 [12] and ZFPM2 [7].

## 4 Discussion

We have introduced a Python-based, parallelized implementation of GeneClass, called PyBoost, with a novel ability to learn regulatory programs according to a specified condition hierarchy, called HiBoost. We include analysis tools for interpretation of the model. Our implementation is an easy-to-install and easy-to-use method using all freely available libraries. As compared to the most popular boosting libraries, such as XGBoost [3], we enable efficient use of paired feature spaces and improved interpretability from a single ADT. Our implementation can be applied to any problem where a 2-dimensional target matrix is predicted from separate featurizations of each dimension of that matrix. We have applied HiBoost to learn interpretable regulatory programs specific to the hematopoietic hierarchy and applied PyBoost to discover novel regulators of tadpole tail regeneration, which were validated experimentally. We hope our implementation will be broadly useful to researchers interested in learning gene regulatory networks in a wide variety of biological contexts.

## Contributions

PG implemented the software, analyzed the hematopoiesis data and wrote the manuscript with feedback from all authors. NH and AK designed the hierarchical algorithm. JC applied the software to regeneration data and experimentally validated predictions. AK provided guidance and feedback.

## Acknowledgements

We would like to thank Nathan Boley and Anna Shcherbina for their help and feedback on the software.

## References

[1] S. Bozdag, A. Li, S. Wuchty, and H. A. Fine. Fastmedusa: a parallelized tool to infer gene regulatory networks. Bioinformatics, 26(14):1792–1793, 2010.

[2] H. J. Bussemaker, H. Li, and E. D. Siggia. Regulatory element detection using correlation with expression. Nature genetics, 27(2):167–174, 2001.

[3] T. Chen and C. Guestrin. Xgboost: A scalable tree boosting system. In Proceedings of the 22Nd ACM SIGKDD International Conference on Knowledge Discovery and Data Mining, KDD ’16, pages 785–794, New York, NY, USA, 2016. ACM.

[4] M. R. Corces, J. D. Buenrostro, B. Wu, P. G. Greenside, S. M. Chan, J. L. Koenig, M. P. Snyder, J. K. Pritchard, A. Kundaje, W. J. Greenleaf, et al. Lineage-specific and single-cell chromatin accessibility charts human hematopoiesis and leukemia evolution. Nature genetics, 2016.

[5] A. Kundaje, S. Lianoglou, X. Li, D. Quigley, M. Arias, C. H. Wiggins, L. Zhang, and C. Leslie. Learning regulatory programs that accurately predict differential expression with medusa. Annals of the New York Academy of Sciences, 1115(1):178–202, 2007.

[6] A. Kundaje, X. Xin, C. Lan, S. Lianoglou, M. Zhou, L. Zhang, and C. Leslie. A predictive model of the oxygen and heme regulatory network in yeast. PLoS Comput Biol, 4(11):e1000224, 2008.

[7] P. Lin, L. J. Medeiros, C. C. Yin, and L. V. Abruzzo. Translocation (3; 8)(q26; q24): a recurrent chromosomal abnormality in myelodysplastic syndrome and acute myeloid leukemia. Cancer genetics and cytogenetics, 166(1):82–85, 2006.

[8] H. Medyouf and J. Ghysdael. The calcineurin/nfat signaling pathway: a novel therapeutic target in leukemia and solid tumors. Cell cycle, 7(3):297–303, 2008.

[9] T. Miyazaki, Y. Kirino, M. Takeno, S. Samukawa, M. Hama, M. Tanaka, S. Yamaji, A. Ueda, N. Tomita, H. Fujita, et al. Expression of heme oxygenase-1 in human leukemic cells and its regulation by transcriptional repressor bach1. Cancer science, 101(6):1409–1416, 2010.

[10] J. J. Moskow, F. Bullrich, K. Huebner, I. O. Daar, and A. M. Buchberg. Meis1, a pbx1-related homeobox gene involved in myeloid leukemia in bxh-2 mice. Molecular and Cellular Biology, 15(10):5434–5443, 1995.

[11] C. G. Mullighan, X. Su, J. Zhang, I. Radtke, L. A. Phillips, C. B. Miller, J. Ma, W. Liu, C. Cheng, B. A. Schulman, et al. Deletion of ikzf1 and prognosis in acute lymphoblastic leukemia. New England Journal of Medicine, 360(5):470–480, 2009.

[12] F. Nakahara, M. Sakata-Yanagimoto, Y. Komeno, N. Kato, T. Uchida, K. Haraguchi, K. Kumano, Y. Harada, H. Harada, J. Kitaura, et al. Hes1 immortalizes committed progenitors and plays a role in blast crisis transition in chronic myelogenous leukemia. Blood, 115(14):2872–2881, 2010.

[13] E. Segal, M. Shapira, A. Regev, D. Pe’er, D. Botstein, D. Koller, and N. Friedman. Module networks: identifying regulatory modules and their condition-specific regulators from gene expression data. Nature genetics, 34(2):166–176, 2003.

[14] C. Zhao, X. Han, Y. H. Zhang, X. Huang, A. Dai, G. Lu, C. C. Yin, L. Chen, and M. J. You. Frequent epigenetic inactivation of msx2 in acute myeloid leukemia. Blood, 116(21):4645–4645, 2010.

